# CD4+ T-cell epitope prediction by combined analysis of antigen conformational flexibility and peptide-MHCII binding affinity

**DOI:** 10.1101/2020.05.21.109967

**Authors:** Tysheena Charles, Daniel L. Moss, Pawan Bhat, Peyton W. Moore, Nicholas A. Kummer, Avik Bhattacharya, Ramgopal R. Mettu, Samuel J. Landry

**Author notes:** Correspondence: Samuel J. Landry.

## Abstract

Antigen processing in the class II MHC pathway depends on conventional proteolytic enzymes, potentially acting on antigens in native-like conformational states. CD4+ epitope dominance arises from a competition between antigen folding, proteolysis, and MHCII binding. Protease-sensitive sites, linear antibody epitopes, and CD4+ T-cell epitopes were mapped in the plague vaccine candidate F1-V to evaluate the various contributions to CD4+ epitope dominance. Using X-ray crystal structures, antigen processing likelihood (APL) predicts CD4+ epitopes with significant accuracy without considering peptide-MHCII binding affinity. The profiles of conformational flexibility derived from the X-ray crystal structures of the F1-V proteins, Caf1 and LcrV, were similar to the biochemical profiles of linear antibody epitope reactivity and protease-sensitivity, suggesting that the role of structure in proteolysis was captured by the analysis of the crystal structures. The patterns of CD4+ T-cell epitope dominance in C57BL/6, CBA, and BALB/c mice were compared to epitope predictions based on APL, peptide binding to MHCII proteins, or both. For a sample of 13 diverse antigens larger than 200 residues, accuracy of epitope prediction by the combination of APL and I-A^b^-MHCII-peptide affinity approached 40%. When MHCII allele specificity is also diverse, such as in human immunity, prediction of dominant epitopes by APL alone approached 40%. Since dominant CD4+ epitopes tend to occur in conformationally stable antigen domains, crystal structures typically are available for analysis by APL; and thus, the requirement for a crystal structure is not a severe limitation.

## Introduction

Rational vaccine design continues to be challenging, due in no small part to the multiple disparate mechanisms and signals that regulate the strength and specificity of the immune response. Antibodies and T cells form the core of adaptive immune responses, but they depend on each other and on potent signals from the innate immune system (1). T-cells recognize polypeptide fragments displayed on the cell surface by polymorphic major histocompatibility complex (MHC) molecules. The two main types of MHC molecules, class I and class II, differ in their source of peptides and the type of type of T cells that recognize them (2, 3). Class I MHC molecules (MHCI) present mostly endogenous peptides and are recognized by CD8+ T cells. Class II MHC molecules (MHCII) present mostly exogenous peptides and are recognized by CD4+ T-cells.

The analysis of large numbers of natural and synthetic MHC-bound peptides, combined with the study of X-ray crystal structures revealed that the specificity of peptide-binding to MHCI and MHCII can be explained by the shape and chemical environment of the peptide-binding cleft (4-6). A substantial degree of variability in peptide specificity derives from the polymorphism of MHCI and MHCII molecules, wherein variant residues in the peptide-binding cleft modulate peptide-binding specificity. In addition, the individual MHCII molecules are substantially more permissive in peptide specificity than MHCI counterparts. Whereas peptide binding to MHCI depends more on contacts with peptide side chains, peptide binding to MHCII depends on hydrogen bonds to the peptide backbone (7). Whereas the MHCI cleft is deep and terminates in closed ends that completely bury the peptide termini, the MHCII cleft is shallow and terminates in open ends that allow the same peptide to bind in multiple different registers. The existence of multiple registers of peptide-binding can explain how a single peptide sequence can give rise to multiple distinct epitopes, and the different registers are thought to be selected by different circumstances of binding (8, 9). Peptide binding is also influenced by the regulated activity of the peptide-MHCII exchange catalyst DM (10).

In general, the MHCI and MHCII display antigens that have been processed in the cytoplasm and endo-lysosome, respectively (2). For MHCI, antigens are targeted for degradation by the ubiquitin proteasome pathway. The antigens are tagged with ubiquitin, unfolded by the ATP-dependent 19S regulatory cap, and then threaded into the proteasome core for degradation. Peptides released from the proteasome are transported by TAP into the endoplasmic reticulum, where they assemble with MHCI during folding. For MHCII, antigens are internalized by pinocytosis, or receptor-mediated endocytosis, or phagocytosis; and proteolyzed by conventional proteases in the endo-lysosome. Then the proteolytic fragments bind to the MHCII, which simultaneously becomes available by the proteolytic processing of its dedicated chaperone, the invariant chain. One notable protein-unfolding activity in the MHCII pathway is the γ-interferon inducible lysosomal thioreductase (GILT). In the absence of GILT, disulfide bonds reshape CD4+ epitope dominance patterns and can severely reduce immunogenicity (11, 12). Major distinctions for the MHCII pathway are the lack of separation between antigen processing and peptide-MHCII binding and the lack of an ATP-dependent unfolding activity in the endo-lysosome, other than acidification.

Current CD4+ epitope prediction tools are based on the binding affinity of the peptide for the MHCII molecule (4). The variability in peptide length, low sequence dependence of peptide binding, and potential for multiple binding registers cause difficulty in predicting MHCII peptide ligands. The potential for proteolytic mechanisms to limit availability of MHCII ligands has long been recognized (13, 14). Recently studies have identified sequence signatures near the ends of MHCII ligands eluted from MHCII complexes (15, 16). These strategies have yielded modest improvements in prediction of MHCII ligands. Although accuracy measured by receiver-operator characteristic (ROC) curves reach 0.75, practical application of prediction has been daunting. When a small, high-scoring fraction of peptides has been tested for restimulation of T-cell responses, a very low hit-rate (< 20%) has been observed (17-20).

Numerous studies have documented a role for antigen conformational stability in antigen processing and epitope presentation (20-24). Studies from this lab have shown that CD4+ T-cell epitopes are found adjacent to flexible regions of the antigen (12, 25-27). These studies led to a generalized model explaining the importance of antigen structure in CD4+ epitope immunodominance (25). In this model, the antigen is proteolyzed within the flexible loops, which allows intervening segments to be separated from the rest of the protein upon binding to the MHCII molecule. The protein segments that are bound to the MHCII molecule continue to be protected while the termini are trimmed by further proteolysis, and then the peptide-MHC complexes are transported to the surface of the cell.

We selected a bacterial subunit vaccine as a model antigen for a study of CD4+ epitope prediction using conformational stability and MHCII binding affinity. Antibodies are crucial for protection against infection by *Yersinia pestis*, causative agent of the Black Plague (28); and antibodies in turn depend on CD4+ T cells for the signals that promote B-cell class switching and affinity maturation. Since the antibodies generally target extracellular proteins, vaccine development has focused on killed organisms, on cellular fractions that include the envelope, cell wall, and capsule, and on secreted protein subunits. Two proteins that have advanced in plague vaccine research are the capsular protein Caf1 and type III secretion component LcrV. In an effort to maximize protectiveness of a single vaccine, genes for Caf1 and LcrV have been fused to produce a single recombinant protein, F1-V (29). Although protective in mice, protection in non-human primates was inadequate; and more advanced vaccine candidates are being developed (30). CD4+ T-cell epitopes for both LcrV and Caf1 have been mapped in mice that had been immunized with the recombinant protein or peptides in multiple mouse strains and using various vaccine formulations (31-33). In the case of Caf1, the efficiency of CD4+ epitope presentation to T-cell hybridomas correlated with availability in the folded structure (34).

Here we have analyzed the potential for native structure in F1-V to shape the pathways of antigen processing, as modeled by F1-V fragmentation in limited proteolysis. Fragmentation was consistent with the accessibility of cleavage sites predicted from the X-ray crystal structures of Caf1 and LcrV and with the accessibility to antibodies against linear epitopes. MHCII-binding and structure-based methods for predicting the CD4+ epitopes of F1-V were evaluated by comparison to epitope maps obtained in three strains of mice. The most accurate epitope prediction arose from a combination of methods that took into account MHC II binding and structure-based limitations on antigen processing.

## Results

### Acid-induced denaturation of LcrV

LcrV was expected to undergo a cooperative unfolding transition as the buffer pH became progressively lower. Acid-induced denaturation of LcrV was monitored by the increase in 4,4’-dianilino-1,1’-binaphthyl-5,5’-disulfonic acid (Bis-ANS) fluorescence, resulting from its increased binding to the denatured protein. Fluorescence was measured with three replicates at each pH. The average fluorescence was calculated from 476 nm to 485 nm for each replicate, then the average and standard deviation were calculated for all replicates at each pH. These values were plotted in an unfolding curve (Fig. 1A). The low level of Bis-ANS fluorescence down to pH 5.6 indicates that F1-V remained at least 92% folded. As the pH was further reduced, LcrV unfolded with the midpoint of the transition at pH 4.7. Limited proteolysis of F1-V with trypsin and proteinase K yielded consistent fragmentation patterns at pH’s as low as 5.6 (Fig. 1B and Fig. S1).

**Figure 1.**
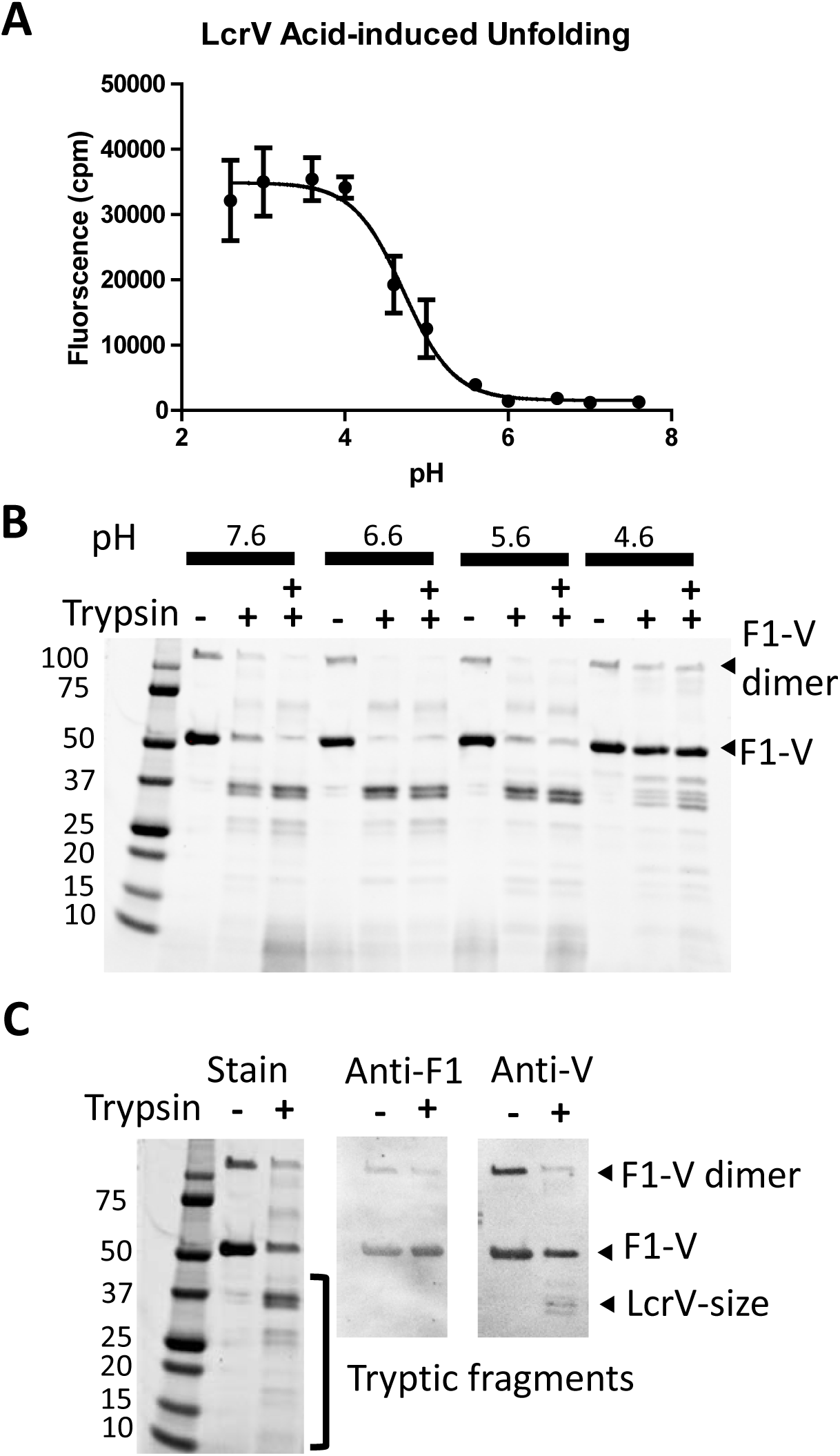
Conformational stability and acid resistance in LcrV and F1-V. A, LcrV resists acid-induced denaturation down to at least pH 5.6, as reported by minimal binding of the fluorescent dye bis-ANS. B, according to SDS-PAGE and Coomassie staining, the fragmentation pattern from limited proteolysis with trypsin remains consistent at pH 7.6, pH 6.6, and pH 5.6, suggesting that the folded structure in F1-V resists proteolysis down to pH 5.6. Two “+” indicates doubling of trypsin concentration. Similar results were obtained for limited proteolysis with proteinase K (Fig. S1). C, in Western blots, prominent 37-kDa fragments from limited proteolysis with trypsin are decorated by anti-LcrV (anti-V) antibodies and thus correspond to LcrV. *BEI*resources depletes the dimeric form of F1-V from the preparation, but an SDS-resistant F1-V dimer constitutes approximately 25% of the mass. The immunological properties of oligomeric and monomeric forms of F1-V were reported to be indistinguishable (68).

### Limited proteolysis of F1-V

We hypothesized that the proteolytic fragmentation of F1-V is primarily controlled by the ability of F1-V segments to conform to the protease active site. Thus, fragmentation patterns from limited proteolysis were expected to reflect the F1-V domain structure and profile of conformational flexibility, and to be somewhat insensitive to the particular protease. For example, the 51 kDa F1-V has 56 potential cleavage sites for trypsin, based on the number of lysine and arginine residues. In spite of the large number of potential cleavage sites, limited proteolysis with trypsin of F1-V yielded a most prominent fragment at 37 kDa, which is the expected size of the C-terminal LcrV portion of F1-V (Fig. 1C). The identity of this fragment as essentially LcrV was supported by Western blotting, which revealed the decoration of an LcrV-sized band with anti-LcrV antibodies but not with anti-Caf1 antibodies. Similar results were obtained from limited proteolysis with other proteases, including elastase, proteinase K, thermolysin, and cathepsin S.

In order to further resolve the identities of proteolytic products and locations of protease-cleavage sites, six major fragments from various protease digestions were excised from the gels and identified by complete tryptic digestion followed by analysis of the peptides with liquid chromatography and mass spectrometry (LC-MS/MS). The 37 kDa fragments from digestion with elastase and cathepsin S were each found to contain tryptic peptides spanning residues 176-479 (Fig. 2A and 3). This segment is approximately 20 residues smaller than expected for the 37 kDa full-length LcrV (171-496). Since our analysis was not optimized for detecting peptides that lack the C-terminal lys/arg or very small peptides (less than eight residues), which would be produced by cleavage in the segment 479-496, we concluded that the 37 kDa fragments corresponded to essentially the full-length LcrV (171-496). LC-MS/MS of the approximately 25-kDa elastase fragment (not shown) and 25-kDa proteinase K fragment (Fig. S1) yielded tryptic peptides that span residues 176-403 and 193-403, respectively, which are consistent with cleavage near the F1-V fusion junction and at a second site C-terminal from residue 403. The 17 kDa elastase fragment yielded tryptic peptides spanning Caf1 residues 75-175. The approximately 30-kDa cathepsin S fragment yielded tryptic peptides that span residues 235-479. The assignment of the 37 kDa, 30 kDa, and 25 kDa fragments to the LcrV portion of F1-V was confirmed by the similar fragmentation pattern obtained from digestion of LcrV by cathepsin S (Fig. 2B).

**Figure 2.**
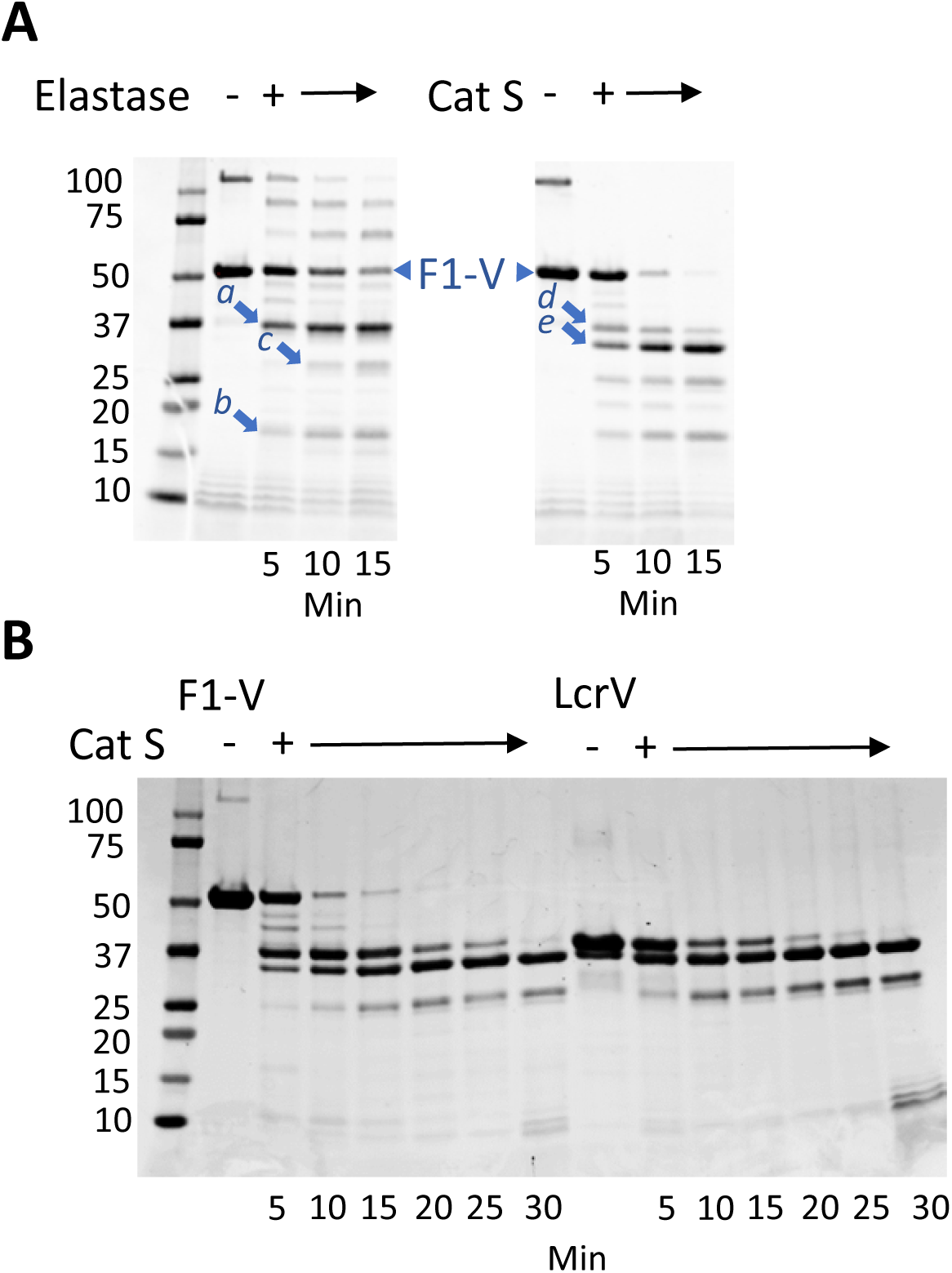
Limited proteolysis of F1-V with elastase and cathepsin S yields similar fragmentation patterns. A, according to SDS-PAGE and Coomassie staining, digestion with elastase and cathepsin S each generate fragments of 37 kDa, 25 kDa, and 17 kDa. Cathepsin S also generates a 30 kDa fragment. Arrows indicate fragments that were identified by tryptic proteomics to contain the following residues of F1-V: fragments *a* and *d*, 176-479; fragment *b*, 75-175; fragment *c*, 176-403; and fragment *e*, 235-479. B, the similarity of fragmentation patterns for digestion of F1-V and LcrV by cathepsin S confirms that the 37 kDa, 30 kDa, and 25 kDa fragments of F1-V correspond to portions of LcrV.

### Peptide-reactive antibody responses

Groups of ten C57BL/6, CBA, and BALB/c mice were immunized intranasally with F1-V combined with the adjuvant, mutant heat-labile toxin (mLT) from *E. coli* (35). Linear antibody epitopes were mapped using the antiserum of each mouse. Of 79 peptides spanning F1-V, ten peptides reacted with antisera from a majority of mice, wherein reactivity was considered positive if the ELISA signal for a peptide exceeded that for the blank by two standard deviations (Fig. 3 and Table S1). These ten peptides occur in four clusters (peptides 18-19, 28-30, 35-36, and 70-72, corresponding to F1-V residues 103-125 (in Caf1), 171-198 (fusion junction), 212-234 (in LcrV), and 422-450 (in LcrV), respectively.

**Figure 3.**
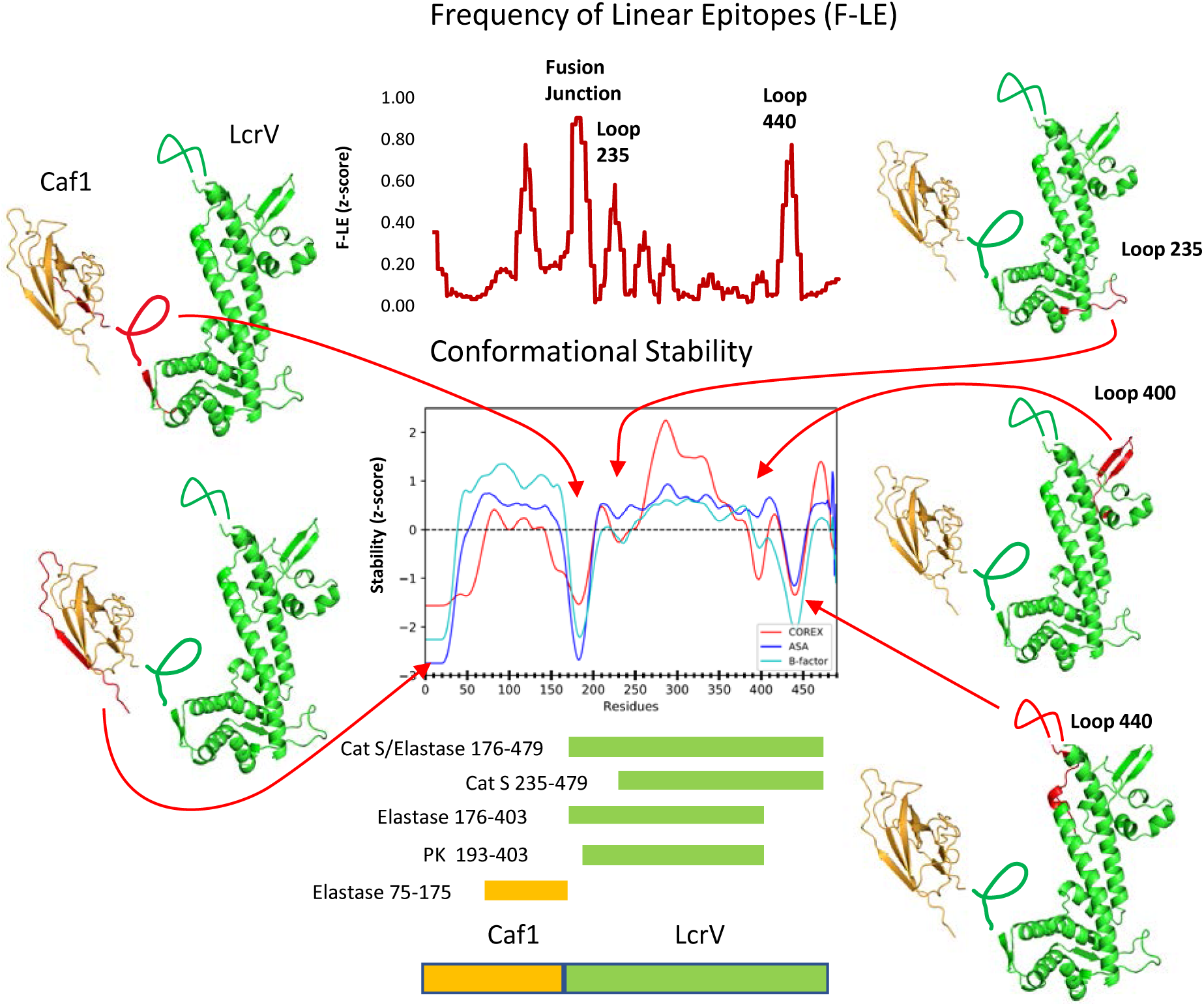
F1-V cleavage sites observed by limited proteolysis correspond to conformationally unstable, solvent-exposed segments. The graph of Conformational Stability illustrates z-score profiles generated from the X-ray crystal structures of Caf1 and LcrV (1Z9S and 4JBU, respectively). The graph of F-LE illustrates a z-score profile of antibody reactivity with linear antibody epitopes. Amber and green bars demarcate proteolytic fragments identified by LC-MS/MS in Caf1 and LcrV, respectively. Preferred sites of proteolytic cleavage are illustrated in the ribbon diagrams, and arrows note their positions in the profiles of conformational stability.

### Proliferative responses

Proliferative responses to F1-V were mapped by peptide restimulation of splenocytes from mice intranasally immunized with F1-V and mLT as described above. Splenocytes from each mouse were tested with individual peptides spanning the complete F1-V. For simplicity, overlapping peptides that could share a single epitope are considered two separate epitopes. Approximately 30% of the peptides were designated “positive” for each group of mice because they stimulated significant proliferation, as scored by the Wilcoxon signed rank test. A large fraction (90%, 71 peptides) were positive in at least one strain. Only a minor fraction (20%) of positive peptides was shared between two or more mouse strains.

Within the Wilcoxon-positive peptide sets, a smaller number of “dominant” peptides stimulated proliferation of splenocytes from at least six mice within a group (Fig. 4 and Table S1). For mouse strains C57BL/6 and CBA, twelve and eleven peptides, respectively, were dominant; and for BALB/c mice, only one peptide was dominant. Approximately one fourth of the 79 peptides (22 peptides) was dominant in at least one strain. Only two peptides were dominant in multiple strains (peptides 13 and 56). Clusters of dominant epitopes were located in three regions of the F1-V: the central stable region of the Caf-1 domain (residues 37-107, peptides 7-17), the central stable region of LcrV (residues 278-348, peptides 46-56), and the C-terminal stable region of LcrV (residues 464-486, peptides 77-78).

**Figure 4.**
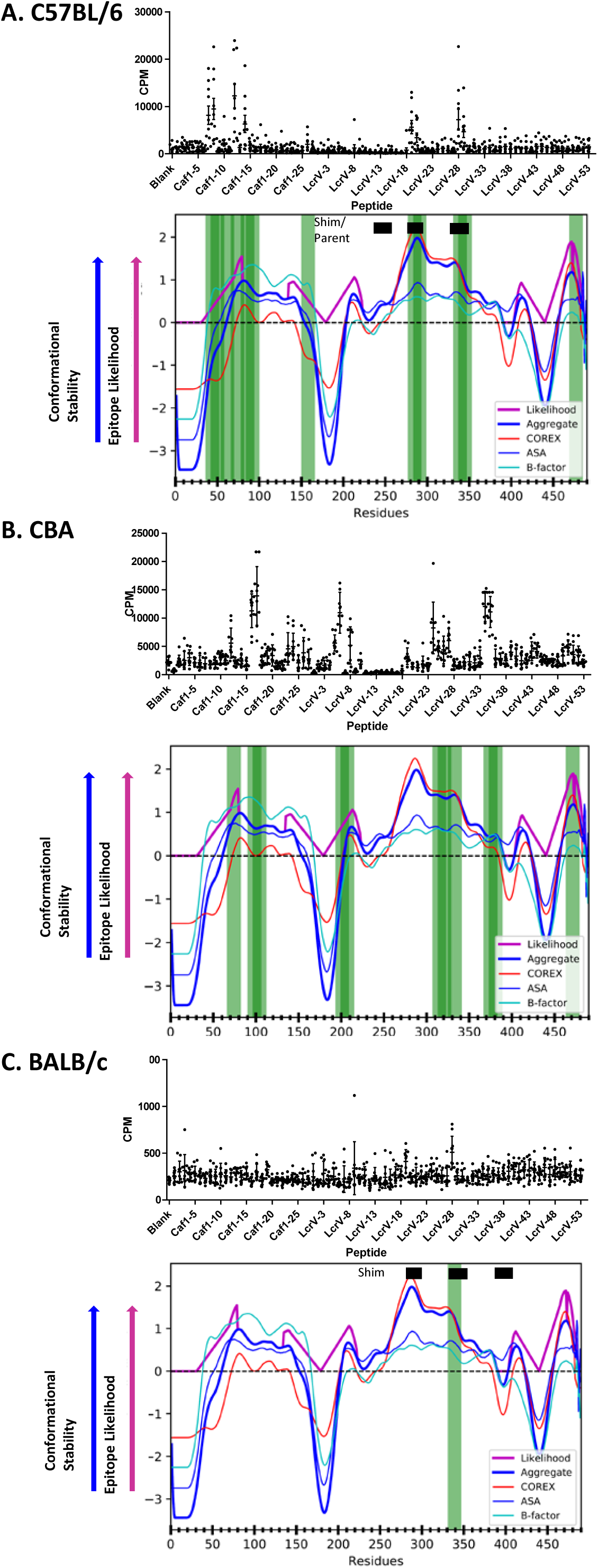
Dominant CD4+ epitopes correspond to stable segments of F1-V. CD4+ T-cell epitopes were mapped in F1-V-immunized mice by splenocyte stimulation with overlapping peptides and measurement of [^3^H]thymidine incorporation, detected as counts per minute (CPM). Vertical green bars indicate dominant epitope-containing peptides, which stimulated proliferation of splenocytes from at least six mice. Z-score profiles for various measures of conformational stability were combined into a single Aggregate z-score. Epitope Likelihood (APL) is a non-zero quantity that tracks with Aggregate conformational stability and is magnified in stable segments adjacent to unstable segments (see Methods). Horizontal black bars indicate T-cell epitopes that were mapped in earlier studies (28,29).

### Accuracy of CD4+ Epitope Prediction

Epitope mapping results were compared to CD4+ epitope predictions based on antigen processing likelihood (APL) and peptide-MHCII binding affinity, as well as a combination of the two approaches. For APL, three measures of residue conformational flexibility/stability (crystallographic b-factor, solvent-accessible surface area, and COREX residue stability) were compiled and then input to the previously described algorithm that assigns APL to stable antigen segments adjacent to flexible antigen segments (36). The algorithm also accepts Shannon sequence entropy as input, but the small number of proteins homologous to Caf1 and LcrV precluded a reliable analysis of sequence entropy; and thus, it was not included. For the MHCII binding analysis, the experimental peptide sequences were entered as input to the web-form for the NETMHCII 2.3 Server; the appropriate H-2 locus was selected; and the resulting values of “1-log50k(aff)” were recorded (4).

As described in the Materials and Methods, we optimized the parameter values for the APL algorithm using a leave-out search over sets of antigens that have been mapped in C57BL/6 mice or in humans. The optimization criterion aims to maximize positive predictive value at the empirical frequency of epitopes observed in the test set. The resulting parameters yield weights on the relative importance of input sources of data as well as for the APL weighting scheme and ratio for combining APL and MHCII scores.

For C57BL/6 datasets (Fig. S2A), the distribution of the data parameter values was on-average 0.13, 0.11, 0.25, 0.50 for sequence conservation, b-factors, COREX and ASA respectively. Notably each of the chosen sources of conformational stability contribute substantially to the optimal PPV results, with COREX and ASA having highest weights, respectively. For the algorithm parameters we obtain an average magnification factor of 1.65 and average loop and flank sizes of 14 and 19 residues, respectively. For the optimal ratio of APL to MHC ratio, we obtain an average of 0.31, and the optimal value of this parameter varies with antigen, ranging from 0.22 for Friend V Env to 0.43 for TMEV Vp2. For datasets from human subjects (Fig. S2B), the distribution of data parameter values was on-average 0.17, 0.18, 0.51 and 0.15 for sequence conservation, b-factors, COREX and ASA respectively. These values are similar to the C57BL/6 dataset, except with COREX and ASA exchanging relative importance. For the algorithm parameters the loop and flank sizes were 16 and 11, while the magnification was optimized at 2.4.

Accuracy of epitope prediction can be assessed in multiple ways. We consider two methods of evaluation, positive-predictive value (PPV) at a particular threshold of prediction score and receiver-operator characteristic area under the curve (ROC-AUC). For the first method we conduct leave-out testing and evaluate the PPV for each antigen at a threshold determined empirically. That is, for each test antigen under consideration, we optimize parameters (see Materials and Methods) and evaluate the number of epitopes identified at a scoring threshold determined by the average frequency of epitopes in the training set used for optimization. For C57BL/6 antigens the empirical frequency was on average 10% (Table 2). We compared the PPV achieved by APL alone, by MHCII-binding alone, and by the Combined scoring methods against a baseline of selecting epitopes at random according to epitope frequency. Neither APL (18%) nor MHCII-binding (27%) achieved significant accuracy, but the PPV for Combined scoring (36%) was significant (Fig. 6A). Our second method of evaluation employed the receiver-operator characteristic (ROC), which offers an overall performance metric that is not tied to a particular threshold of prediction. The ROC curve for each prediction can be evaluated by the area under the curve (AUC) of sensitivity versus 1-minus-specificity (false-positive level). If the AUC exceeds a value of 0.5 (p < 0.05), then the scores can be considered to have predictive power better than random. For a peptide collection corresponding to the C57BL/6 epitope-mapping studies (600 peptides), the ROC-AUC achieves significance at a value of approximately 0.57. By this metric, APL and MHCII-binding each have predictive power, and Combined is best (Fig. 6B).

**Table 2.** Number of peptides: epitopes, highly-scoring, and correctly predicted for C57BL/6 mice

| Antigen | Epitopes | Ref. | PDB ID | Total<br>Peptides | Threshold<br>(No. of Peptides) | Correctly Predicted |  |  |
| --- | --- | --- | --- | --- | --- | --- | --- | --- |
|  |  |  |  |  |  | APL | MHCII<br>Binding | Combined |
| HIV gp120 | 5 | (12) | 3JWO | 46 | 0.10 (4) | 0 | 1 | 1 |
| Plague F1-V | 12 | This work | 1P5V,4JBU | 78 | 0.09 (7) | 2 | 2 | 2 |
| M.t. Ag85A | 5 | (46) | 1DQZ<sup>1</sup> | 27 | 0.09 (2) | 0 | 0 | 0 |
| Friend V. Env | 6 | (47) | 1AOL | 37 | 0.09 (3) | 1 | 2 | 2 |
| GFP | 1 | (48) | 2QLE | 17 | 0.10 (1) | 1 | 0 | 1 |
| *M.t.* Mpt51 | 1 | (49) | 1R88 | 25 | 0.10 (2) | 0 | 0 | 0 |
| *M.t.* PstS | 3 | (50) | 1PC3 | 32 | 0.10 (3) | 0 | 1 | 1 |
| VSV-G | 2 | (51) | 2CMZ | 44 | 0.10 (4) | 0 | 1 | 1 |
| OVA | 9 | (52) | 1OVA | 75 | 0.10 (7) | 0 | 3 | 3 |
| Flu HA | 9 | (53) | 3LZG | 83 | 0.09 (7) | 2 | 1 | 2 |
| Flu NA | 9 | (54) | 3CYE | 63 | 0.09 (5) | 1 | 2 | 2 |
| LLO | 3 | (55) | 4CDB | 48 | 0.10 (4) | 1 | 1 | 1 |
| TMEV VP2 | 1 | (56) | 1TME | 25 | 0.10 (2) | 0 | 1 | 1 |

<sup>1</sup>homology model template

For antigens mapped in human subjects, we can evaluate only APL scoring predictions because information about the restricting MHCII alleles is incomplete. Accuracy for APL scoring at the empirical thresholds (top 17%, on average) was not significantly greater than random sampling. However, when we loosen the threshold to provide 50% more peptides than the epitope frequency (top 25%, on average), we achieve a significant PPV (27%). Further loosening the threshold does not improve PPV, presumably because adding more peptides does not capture enough true-positives to outweigh the additional false-negatives. The ROC-AUC of 0.71 is highly significant.

We sought to improve the accuracy of APL in predicting F1-V epitopes in by supplementing or replacing the crystallographic data with biochemical evidence of conformational flexibility and protease sensitivity. The frequency of linear antibody epitopes (F-LE) was converted to a residue-by-residue z-score that could be included alongside B-factor, solvent-accessible surface area, and COREX residue stability. Likewise, protease sensitivity/resistance (as a binary score) was assigned to all residues that were excluded from proteolytic fragments and to the residues at the termini of fragments (Fig. 3). The supplementation of APL-input with these profiles did not substantially change the results, suggesting that the crystallographic and biochemical approaches yield similar information about the F1-V structure with regard to epitope prediction (Table 1). Remarkably, the combination of the two biochemical parameters F-LE and protease sensitivity/resistance (without the biophysical parameters) achieved significant accuracy in the prediction of F1-V CD4+ epitopes in CBA mice (Fig. 6 and Table 1).

**Table 1.** Accuracy of prediction methods for F1-V CD4+ epitopes

| Prediction Method | Hits for 7 peptides predicted at the Top<br>10<sup>th</sup> percentile and ROC-AUC |  |
| --- | --- | --- |
|  | C57BL/6 | CBA |
| APL | 2 / 0.72* | 1 / 0.69* |
| MHCII | 2 / 0.76** | 2 / 0.53 |
| APL + MHCII | 2 / 0.82*** | 1 / 0.62 |
| APL (w/F-LE) | 2 / 0.73* | 3 / 0.69* |
| APL (w/F-LE) + MHCII | 4 / 0.81*** | 2 / 0.64 |
| APL (w/proteolysis) | 2 / 0.69* | 3 / 0.69* |
| APL (w/proteolysis) + MHCII | 4 / 0.79** | 3 / 0.63 |
| APL (F-LE only) | 2 / 0.57 | 1 / 0.64 |
| APL (Proteolysis only) | 3 / 0.63 | 1 / 0.73* |
| APL (F-LE + proteolysis) | 0 / 0.56 | 3 / 0.75** |
| APL (F-LE + proteolysis) + MHCII | 3 / 0.71* | 3 / 0.69* |

\*, p < 0.05; \*\*, p < 0.01; \*\*\*, p < 0.001

## Discussion

Protease-sensitive sites and CD4+ T-cell epitopes were mapped in F1-V in order to investigate the influence of antigen processing on epitope dominance. Antigen processing for the MHCII pathway occurs in an endo-lysosome-like compartment, where antigens, proteases, the disulfide-exchange catalyst GILT, MHC II, and the MHCII-peptide-exchange catalyst DM together experience a time- and activation-dependent acidification. Since many proteins retain native-like conformations in acidic environments, we sought to test the hypothesis that F1-V three-dimensional structure influences CD4+ epitope dominance by limiting access to lysosomal proteases and the MHCII molecule.

We hypothesized that conformationally stable domains of F1-V could resist acidification in the antigen-processing compartment. LcrV resisted denaturation to at least pH 5.6, where 92% of the protein remained in the native structure, as reported by Bis-ANS binding (Fig. 1). The acid-induced unfolding of the complete F1-V was not undertaken because the complex unfolding pathway of the multidomain protein would preclude analysis of the fraction unfolded. As a separate molecule, Caf1 adopts an incompletely folded metastable state prior to assembly into fibers, and its structure has only been characterized in complexes with the dedicated molecular chaperone Caf1M (37). An N-terminal segment of the protein must either associate with the chaperone or fold with another subunit of Caf1. Recombinant Caf1 forms a flocculant aggregate that resembles the capsule formed by *Yersinia pestis*, and denaturation reduced its protectiveness as a vaccine (38). Presentation of Caf1 epitopes to T-cell hybridomas was found to be increasingly dependent on antigen processing according to their position from N-terminus to C-terminus, suggesting that the structure was progressively unraveled during processing (34). Findings here indicate that LcrV should be added to a list of antigens, including Caf1, tetanus toxoid, Bet v 1, RNAse A, horseradish peroxidase, and *P. auruginosa* exotoxin domain III, for which the conformations have been shown to resist acid denaturation sufficiently to limit access to proteolytic enzymes and/or MHCII molecules (21, 23, 24, 38, 39).

Limited proteolysis and peptide-mapping of F1-V indicated that the most proteolytically sensitive sites lie near the Caf1-LcrV fusion junction. A prominent fragment of size equal to LcrV (37 kDa) and that reacted with anti-LcrV antibodies was generated by each of the tested proteases (trypsin, protease K, elastase, thermolysin, and cathepsin S). Representative 37 kDa fragments from digestion with elastase and proteinase K were found by tryptic proteomics to contain the segments 176-479 and 193-479, respectively (Fig. 3). For comparison, in denaturing conditions, the exhaustive fragmentation of LcrV with trypsin yielded 28 peptides, none of which was larger than 4 kDa (data not shown). Thus, preferred cleavage of F1-V by trypsin and other proteases near the F1-V fusion junction was most likely due to conformational disorder in the region of the fusion junction; whereas, the Caf1 and LcrV portions of F1-V remained relatively ordered and resistant to proteolysis.

Proteolytically sensitive sites within the LcrV portion of F1-V are associated with conformationally disordered loops that are evident in the LcrV crystal structure. A 30 kDa cathepsin-S fragment of LcrV spanning residues 235-479 resulted from cleavage within a flexible loop (Loop-235) that protrudes from the N-terminal globular domain of LcrV (Fig. 3). Preferred C-terminal cleavage sites in F1-V were located within the C-terminal twenty residues (479-496) of LcrV or within a large unstable region of LcrV spanning residues 390-460, including hairpin Loop-400 and disordered Loop-440. Early fragments are likely to be subject to rapid further proteolytic cleavage, as the result of enhanced conformational flexibility in the new terminal regions. For example, we suspect that initial cleavage occurs in the large flexible Loop 440 and that additional proteolytic steps shorten the polypeptide from the C-terminus to a point between residues 403 and 409, based on the tryptic proteomics.

A residue-level profile of linear antibody epitopes potentially offers an alternative source of conformational stability data that can supplement or replace crystallographic data. Antiserum reactivity with a synthetic peptide suggests that the antibody epitope is contained within the corresponding F1-V segment and that the epitope is available in the context of the native protein. In order to be available, the segment must be solvent-accessible and able to conform to the binding site on the antibody (40, 41). The majority of peptides that reacted with antibodies raised against the intact antigen corresponded to conformationally unstable antigen segments (Fig. 3). The frequency of linear epitopes (F-LE) was scored as the average fraction of mice that reacted with the peptides that contain the residue. Each residue appears in three peptides, and each peptide was probed with immune serum of ten mice from each of three mouse strains. Three clusters of the most consistently reactive peptides were in segments that were recognized as disordered using biophysical data, one near the fusion junction and two internal loops of LcrV (Loop-235 and Loop-440).

Proliferative T-cell responses were well distributed in the F1-V sequences and punctuated by dominant epitopes that obtained responses in a majority of mice of a given strain. For the two mouse strains (C57BL/6 and CBA) having a single MHCII molecule (I-A^b^ and I-A^k^, respectively), approximately half of the peptides that scored positive using the Wilcoxon Signed Rank Test obtained a significant response in a majority of mice – and therefore are also defined as dominant epitopes. Only one peptide was dominant in BALB/c mice. While only two of 22 dominant epitopes were shared between strains, clusters of dominant epitopes were shared among strains, e.g., spanning residues 79-113 or 330-355 (Fig. 4). The clustering of non-identical epitopes could be explained by distinct but overlapping MHCII sequence preferences or by the limitation of MHCII selection to antigen segments that preferentially emerge from antigen processing.

Two of three LcrV epitopes that were previously identified in C57BL/6 mice were re-identified here, and one of three LcrV epitopes previously identified in BALB/c mice was re-identified here (Fig. 4). Among the possible explanations for the differences are the use of F1-V instead of LcrV as the immunogen and different routes and adjuvants for the immunization (31, 32). Our studies of limited proteolysis yielded no evidence that fusion to Caf1 had affected the conformation of LcrV. Thus, we favor the conclusion that route and/or adjuvant affected antigen processing through engagement of different antigen presenting cells or modulation of the agents of antigen processing.

Essentially all of the dominant epitopes occurred within conformationally stable segments of F1-V, as represented by a positive Aggregate z-score (Fig. 4). However, the dominant epitopes were not centered on the stable segments. Rather they appear on the edges of the stable segments, as represented by the epitopes at residues 65-80 in both C57BL/6 and CBA mice and at residues 330-355 in all three strains of mice. This offset from center of stability was the basis for development of the antigen processing likelihood (APL) algorithm, which upweights the prediction on the edge of stable segments (36).

Selected dominant epitopes reveal strengths and weaknesses of APL and MHCII-binding for epitope prediction. An important strength of APL is its potential for eliminating false-positives from the MHCII(I-A^b^) binding profile. A prominent example for F1-V in C57BL/6 mice in is noted for peptides 2-5 C57BL/6 mice. All four peptides are predicted to be epitopes by MHCII(I-A^b^) binding (Fig 5A, circle). Although peptides 2-4 obtained responses in 4 mice, none of these peptides was dominant; and this can be explained by their location within the unstable N-terminal segment (residues 19-35) of Caf1 (Figs. 3 and 4). The poor immunogenicity of peptides 2-5 is most likely due to destructive processing, as reflected in the low APL scores. In contrast, the nearby dominant peptides 8 and 12 scored highly in both MHCII binding and APL. APL also has the potential to correct false-negatives. For example, moderate MHCII(I-A^b^) binding at peptides 46 and 47 was boosted by APL into the top 11^th^ percentile of the Combined prediction (Fig. 5A, 5B, square). In all of these examples, we surmise that antigen-processing modulates the effective concentration of MHCII-binding sequences.

**Figure 5.**
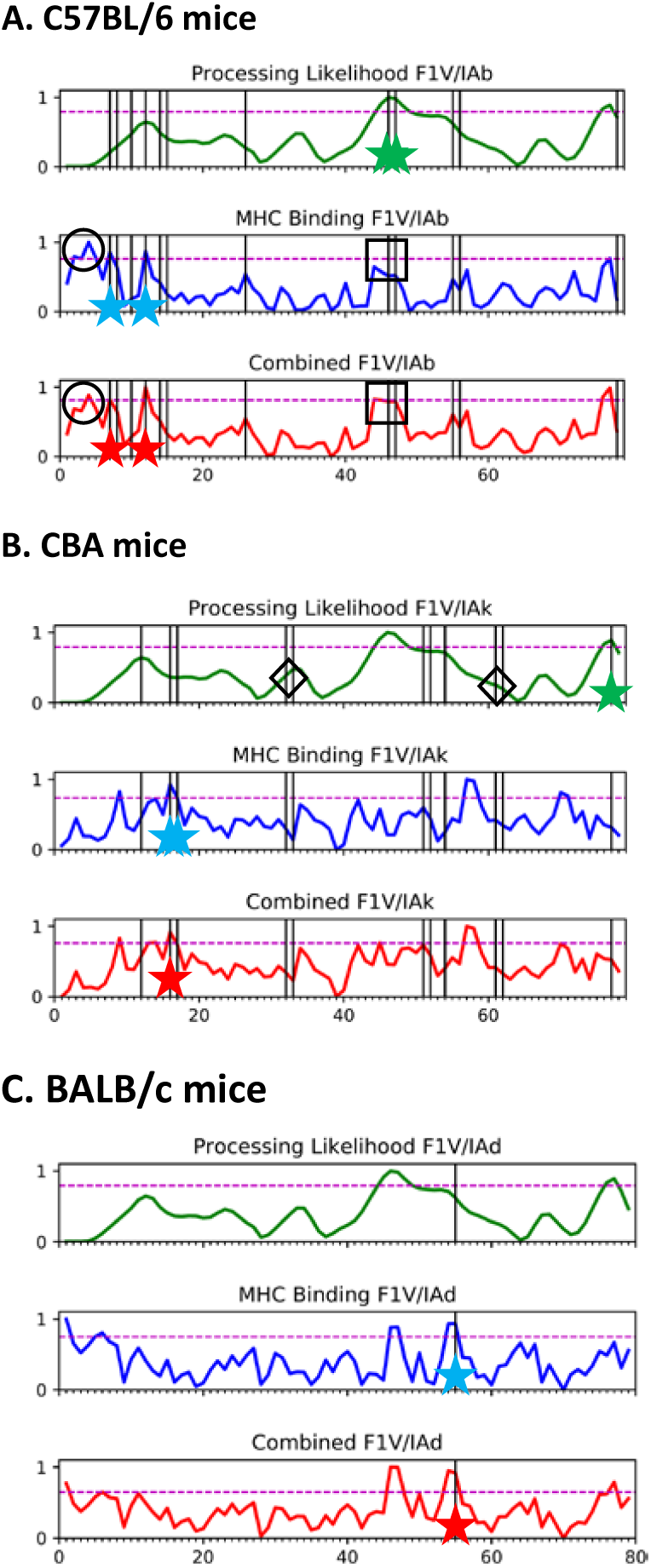
The combination of APL and MHCII-binding performs better than MHCII-binding alone for prediction of dominant CD4+ epitopes of F1-V in C57BL/6 and CBA mice. Dashed lines indicate the 85^th^ percentile of prediction score. Vertical lines indicate dominant epitopes identified by T-cell proliferation. A larger number of dominant epitopes was correctly predicted by the combination of APL and MHCII binding (stars). Certain false-positives in the MHCII-binding prediction (open circles) were eliminated when combined with the APL prediction. Some false-negatives in the APL prediction might be corrected by improvement of the algorithm (open diamonds).

**Figure 6.**
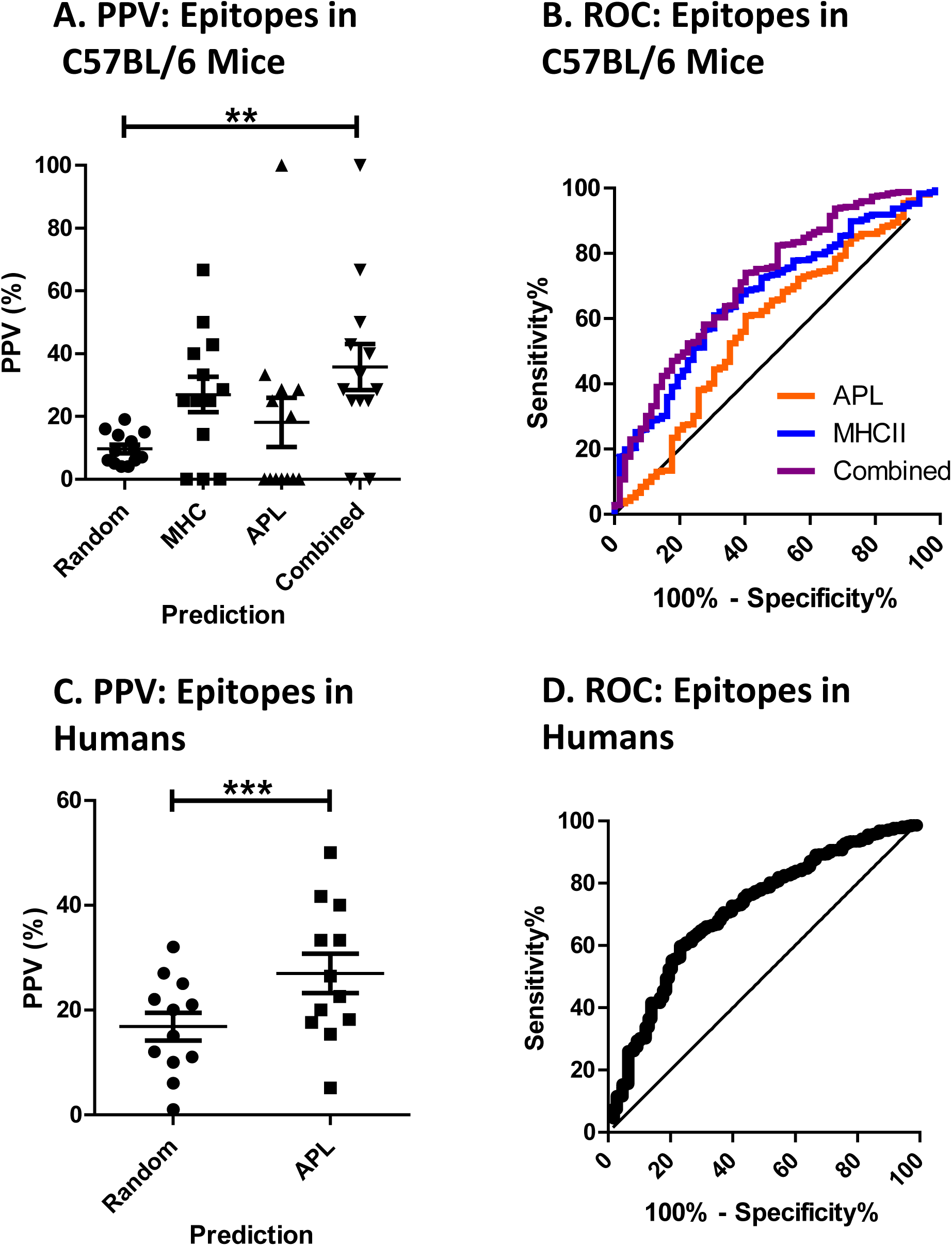
Accuracy of CD4+ epitope predictions for antigen collections. A, accuracy for thirteen antigens in C57BL/6 mice, as illustrated by ROC curve. The indicated AUC values are significantly greater than 0.5 (p < 0.0001). The inset highlights the difference in accuracy between prediction methods for the region of high sensitivity. B, Frequency of epitope-hits among peptides scoring above threshold (approximately top 10%) peptides predicted for individual antigens using the indicated method. “Random” indicates the frequency of experimentally observed epitopes within the complete set of peptides for the antigen (i.e., frequency of epitope-hits for random sampling). *Asterisks* indicate significance by One-way ANOVA with repeated measures and Tukey method of multiple comparisons (*, p < 0.05; **, p < 0.01). C, accuracy for twelve antigens in humans, as illustrated by ROC curve. The indicated AUC value is significantly greater than 0.5 (p < 0.0001). D, Frequency of epitope-hits among peptides above 1.5 x threshold (approximately top 17%) scored for individual antigens using the indicated method. The *asterisk* indicates significance by paired T-test, p < 0.05.

An important weakness in APL may be represented by its failure to predict dominant epitopes in peptides 51-52 and 61-62 in CBA mice (Fig. 5B, diamonds). Peptides 51-52 lie adjacent to the highly flexible and protease-sensitive fusion junction. Although we might expect these peptides to be well processed, they did not score in the top 10^th^ percentile of APL because this segment of F1-V is not as conformationally stable as other segments of the protein (Fig. 4B, note lower stability of residues 200-220 and 370-390, compared with other immunogenic segments). The APL algorithm assigns a score to residues according to conformational stability and upweights the score of residues adjacent to unstable segments in proportion to the conformational stability of the residue receiving the score. Due to modest local stability, peptides 51-52 were not upweighted into the top 10^th^ percentile of APL score. Likewise, peptides 61 and 62 were not upweighted into the top 10^th^ percentile even though they are on the N-terminal flank of Loop-400. Strategies to overcome this weakness in APL scoring are under investigation. Remarkably, APL based solely on the biochemical analysis (F-LE + proteolysis) predicted both the 51-52 and 61-62 dominant epitopes (Fig. 6). Thus, efforts to improve the prediction of proteolytic sensitivity from crystallographic data may also yield an improvement in epitope prediction.

For a collection of thirteen antigens whose CD4+ T-cell epitopes have been systematically mapped in C57BL/6 mice, the combination of APL and MHCII(I-A^b^) surpassed either prediction method alone (Table 2). For each antigen, mice were primed with the antigen in a native-like state (e.g., not by peptides) and then probed by lymphocyte restimulation using a complete set of overlapping peptides spanning the antigen. The set contains no antigens of less than 200 amino acid residues and no more than one antigen of a homologous protein family (e.g., flavivirus envelope proteins). The size-minimum accounts for the fact that small proteins tend to have reduced conformational stability and lack complex domain structure (42). Thus, small proteins yield little complexity in fragmentation by limited proteolysis, and the structures have little effect on antigen processing and epitope dominance. For this group of model antigens, an average of ten percent of the tested peptides contained epitopes. As described in the Materials and Methods, we evaluated the positive predictive value (PPV) for each antigen at a scoring threshold determined by remaining antigens, on average the top 10^th^ percentile. In a real-life test, this is comparable to choosing seven of seventy-eight possible peptides from plague F1-V. For this protein, MHCII(I-A^b^) binding, APL, and the Combined prediction each correctly scored two hits (Table 2). For the thirteen antigens, the average accuracy (using the antigen-specific thresholds) was 18% for APL, 27% for MHCII(I-A^b^) binding, and 36% for the Combined prediction, and only the Combined method achieved significant accuracy when the three methods were tested side-by-side (Fig. 7A). The Combined prediction also delivered at least one hit in all but two of the thirteen antigens (Table 2).

**Figure 7.**
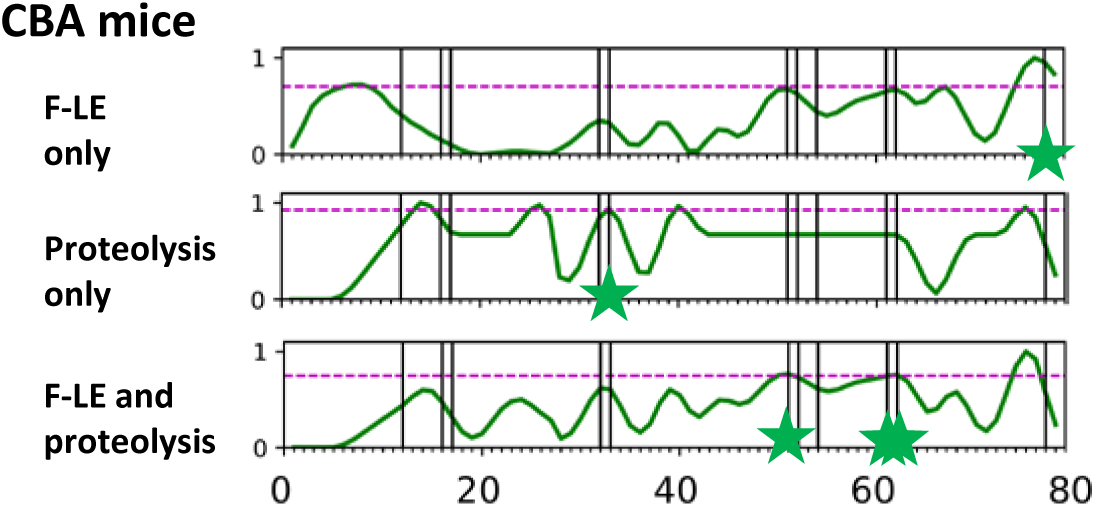
Prediction of dominant CD4+ epitopes of F1-V by APL without crystallographic structures. Z-score profiles generated on the basis of F-LE (Fig. 3) and fragmentation by limited proteolysis (see text) were used as input to the APL algorithm. APL using only these two parameters identified a significant number of the dominant F1-V epitopes in CBA mice (stars).

APL could be a major asset for the prediction of CD4+ epitopes in human immune responses. In the original description of the algorithm, APL correctly identified epitopes at a rate of 23% of peptides in the top-20^th^ percentile for a set of nine systematically mapped antigens (36). Here, we have updated the set to eliminate antigens of less than 200 residues, include three new antigens, and replaced antigen-85A with its homolog, antigen-85B (Table 2). Antigen-85B was considered superior because its dominant epitopes were characterized by frequency in the cohort of human subjects, rather than average intensity, i.e., number of Elispots. For the updated set of antigens, the experimentally mapped epitopes occurred with a frequency of 17% within the series of overlapping peptides spanning the antigens. For thresholds 50% higher than the experimental frequency, the hit-rate was 27%, and at least one epitope was correctly identified for all twelve antigens (Fig. 7C and Table 3). These results for APL are comparable to the best CD4+ epitope predictions so far reported for human epitope-mapping data (16, 43).

**Table 3.** Number of peptides: epitopes, total, and correctly predicted for human subjects

| Antigen | Epitopes | Ref. | PDB ID | Total<br>Peptides | Threshold<br>(No. of Peptides) | APL |
| --- | --- | --- | --- | --- | --- | --- |
| Ad5 hexon | 16 | (57) | 3TG7 | 134 | 0.17 (23) | 8 |
| *M.t.* Ag85b | 9 | (58) | 1F0N | 28 | 0.16 (4) | 1 |
| *M.l.* Hsp70 | 11 | (59) | 2V7Y<sup>2</sup> | 49 | 0.16 (8) | 2 |
| NS3 helicase | 7 | (60) | 1CU1 | 45 | 0.17 (7) | 1 |
| Polio Vp1 | 5 | (61) | 1VBC | 24 | 0.16 (3) | 0 |
| TBE Env | 12 | (62) | 1URZ | 120 | 0.18 (21) | 2 |
| Tetanus toxoid | 10 | (63) | 1Z7H | 51 | 0.17 (8) | 3 |
| Ves v 5 | 7 | (64) | 1QNX | 65 | 0.18 (11) | 1 |
| PE38-III | 17 | (48) | 1IKQ | 67 | 0.16 (10) | 6 |
| Flu HA | 6 | (65) | 2VIU,1HTM | 98 | 0.18 (17) | 3 |
| HIV GAG | 6 | (66) | 4XFX<sup>1</sup> | 22 | 0.16 (3) | 1 |
| *M.t.* Mal6G | 2 | (67) | 6DNP | 143 | 0.18 (26) | 2 |

<sup>1</sup>homology model template

An important limitation of APL is the requirement of a high-resolution three-dimensional structure, typically an X-ray crystal structure. In a survey of the protein universe, 70% of sequences can be at least partially modeled with an existing experimental structure (44). For the purposes of CD4+ epitope prediction, such homology-modeled structures are adequate. Moreover, many of the protein sequences that cannot be modeled are likely to be natively unfolded and therefore poorly immunogenic for CD4+ T cells because they are protease-sensitive (23, 41). Thus, APL may prove to be more useful than the structure-requirement might initially suggest.

## Materials and Methods

### Proteins and Peptides

Recombinant subunit vaccines LcrV (also known as V-antigen) and F1-V, were obtained from Biodefense and Emerging Infections Research Resources Repositiory (*BEI*resources). The 17-mer peptides spanning the entire F1-V were also obtained from *BEI*resources. The peptides spanned Caf1 in 6-residue steps and overlapped by 11 residues and spanned LcrV in 5-residue steps and overlapped by 12 residues. Peptides were dissolved in 1 mL of dimethyl sulfoxide (DMSO), 0.05 % trifluoroacetic acid, 70% acetonitrile, or 6M guanidine-HCl as recommended by the supplier to obtain a final concentration of 1 mg/mL.

### Acid-induced denaturation of LcrV

For acid induced unfolding experiments the hydrophobic dye BIS-ANS (4,4’-dianilino1,1’-binaphthyl-5,5’-disulfonic acid, Invitrogen) was used to monitor protein unfolding by fluorescence spectroscopy with an excitation wavelength of 390 nm. Emissions were scanned from 400 nm to 500 nm with a Photon Technology International Fluorescence Spectrometer. Different pH conditions were generated using phosphate-citrate buffer, where 0.2 M dibasic sodium phosphate and 0.1 M citric acid were mixed until the desired pH was reached. Protein was mixed with dye in phosphate-citrate buffer ranging from pH 7.6 to 2.6 at a concentration of 0.1 µM protein and 1.0 µM dye. Fluorescence intensities averaged for the range 476 nm to 485 nm and for three replicates were analyzed by nonlinear regression using a sigmoidal dose-response variable slope regression model (Prism 6).

### Limited Proteolysis

Proteolysis experiments were conducted in phosphate-citrate buffer at the indicated pH and formulated as above. After incubation all proteolysis reactions were terminated by the addition of an equal volume of BioRad Laemmli Sample Buffer containing 1 mM phenylmethylsulfonyl fluoride (PMSF) and 150 mM 2-mercaptoethanol. Samples were boiled for 5 minutes and then analyzed by sodium dodecyl sulfate-polyacrylamide gel electrophoresis (SDS-PAGE), using a 4 to 20% BioRad TGX gradient gel. Gels were stained with Coomassie blue (45) and scanned on a BioRad Chemidoc MP imaging system and analyzed with BioRad ImageLab software. Trypsin limited proteolysis experiments were performed in a volume of 20 µL with 5 µg of F1-V protein and 0 µg, 1.25 µg or 2.50 µg of trypsin for 30 minutes at 25 °C. Elastase limited proteolysis experiments were performed in a volume of 300 µL phosphate-citrate buffer pH 7.6 with 90 µg F1-V protein and 0.9 µg of Porcine Elastase (Milipore-Sigma). The reaction mixture was incubated at 37 °C for 1 hour with 20 µL aliquots taken every 5 minutes and dispensed into microcentrifuge tubes containing loading buffer, boiled and analyzed as described above. Cathespin S limited proteolysis experiments were performed in a volume of 40 µL phosphate-citrate buffer pH 7.6 containing 18 µg F1-V protein or LcrV protein, 5 mM dithiothreitol, and 0.5 µg Cathepsin S (Milipore-Sigma). The reaction mixture was incubated at 37 °C for 30 minutes with 5 µL aliquots taken every 5 minutes. Aliquots were prepared for and analyzed by SDS-PAGE as described above. Thermolysin limited proteolysis experiments were performed in 100 mM sodium acetate buffer pH 5.5 containing 10 mM CaCl_2_ and 50 mM NaCl. 5.0 µg of F1-V protein was digested with 0.025 µg, 0.0025 µg or 0.00025 µg of thermolysin for 20 minutes at 25 °C. Reactions were stopped by the addition of loading buffer also containing 100 mM ethylenediaminetetraacetic acid (EDTA) and analyzed by SDS-PAGE as described.

### Mass Spectrometry of Limited Proteolysis Fragments

Proteolytic cleavage sites were identified by trypsin sequencing of fragments excised from SDS-PAGE gels. Briefly, each gel slice was destained using a 20-volume excess of 50 mM ammonium bicarbonate and 50% methanol for 20 minutes, twice. Destained gel slices were dehydrated by incubating in a 20 volume excess of 75% acetonitrile for 20 minutes. Dried slices were then incubated in a 5-volume excess of 20 µg/ml mass-spectrometry-grade trypsin dissolved in 50 mM ammonium bicarbonate at 37 °C overnight. Each sample was subjected to a 60-minute chromatographic method employing a gradient from 2-25% acetonitrile in 0.1% formic acid (ACN/FA) over the course of 30 minutes, a gradient to 50% ACN/FA for an additional 10 minutes, a step to 90% ACN/FA for 8 minutes and a re-equilibration into 2% ACN/FA. Chromatography was carried out in a trap-and-load format using a PicoChip source (New Objective, Woburn, MA); trap column C18 PepMap 100, 5 µm, 100 Å and the separation column was PicoChip REPROSIL-Pur C18-AQ, 3 µm, 120 Å, 105 mm. The entire run was µl/min flow rate. Survey scans were performed in the Orbitrap utilizing a resolution of 120,000 between 375-1600 m/z. Data dependent MS2 scans were performed in the linear ion trap using a collision induced dissociation (CID) of 25%. Raw data were searched using Proteome Discoverer 2.2 using SEQUEST. The Protein FASTA database was *M. musculus* (TaxID = 10090) version 2017-07-05 with the PE-III sequence added. Static modifications included carbamidomethyl on cysteines (= 57.021) and dynamic modification of oxidation of methionine (= 15.9949). Parent ion tolerance was 10 ppm, fragment mass tolerance was 0.6 Da, and the maximum number of missed cleavages was set to 2. Only high scoring peptides were considered utilizing a false discovery rate (FDR) of 1%.

### Western Blots of Proteolysis Experiments

After limited proteolysis of F1-V with trypsin SDS-PAGE gels were run as described. Following electrophoresis, gels were transferred to a polyvinylidene difluoride membrane using the Bio-Rad transblot turbo transfer system and packs. Membranes were briefly stained with ponceau (Milipore-Sigma) and imaged. Membranes were then blocked for 1 hour at room temperature with phosphate buffered saline and 0.05% Tween-20 (PBST) containing 2.5% non-fat dry milk (NFDM). Membranes were rinsed once with PBST and incubated overnight at 4 °C with polyclonal goat anti-serum specific for F1-V or LcrV (BEI resources #NR-31024 and #NR-31022) diluted 1:10,000 in PBST plus 2.5% NFDM. The following day blots were washed 3 times with PBST for 5 minutes at room temperature followed by a 1-hour incubation with the donkey anti-goat Alexa flour 488 conjugated secondary antibody (Invitrogen) at a 1:10,000 dilution in PBST plus 2.5% NFDM in the dark. Blots were washed 3 times for 5 minutes with PBST and imaged on a Bio-Rad Chemidoc MP imaging system.

### Immunization

Groups of ten 6- to 8-week-old female C57BL/6 and CBA/J mice were obtained from Jackson Laboratories and ten 6- to 8-week-old BALB/c mice were obtained from Charles River Laboratories, Inc. The mice were immunized intranasally with 10 µg of recombinant subunit vaccine, F1-V and 5 µg of mutant (R192G) heat-labile toxin (mLT) as adjuvant in a final volume of 10 µL. During immunizations mice were anesthetized using isoflurane in O_2_ within an induction chamber. Intranasal administration was delivered by pipetting 5 µL into each nostril. The mice received two boosts of the same mixture at two-week intervals. Mice were sacrificed via CO_2_ asphyxiation one week after the final boost. Spleens and cardiac blood were obtained from each mouse and processed further. All mouse experiments followed institutional guidelines approved by the Tulane Animal Care and Use Committee.

### Isolation of mouse serum

Immediately following CO_2_ asphyxiation, cardiac blood was drawn using a 1 ml syringe with a 21-gauge needle. Whole blood was placed in a BD Microtainer gold serum tube (BD Biosciences) for at least 1 hour before processing. To collect serum, tubes were centrifuged for 90 seconds at 13,000 rpm in a table-top centrifuge. The serum from each mouse was transferred into sterile 1.5 ml microcentrifuge tubes and stored at – 20 °C.

### Mapping of linear B-cell epitopes

A single peptide at a concentration of 4 µg/ml, in 0.1M sodium bicarbonate, was added to each well of a flat-bottom 96-well plate and placed at 4 °C overnight. The next morning, the plates were washed 5 times with ELISA wash buffer (10% Triton-X in PBS), using an ELISA plate washer, and blocked in PBS containing 200 µL of 0.5% Tween20, 4% whey, and 10% fetal bovine serum (PBS/T/W + 10% FBS) for 30 minutes at room temperature. Blocking buffer was removed, and the plate was washed. The mouse serum (primary antibody) was diluted in PBS/T/W + 10% FBS to a final concentration of 1:100 and 100 µL was placed on the plate for 1 hour at room temperature. After 1 hour, the plates were washed and 100 µL of secondary antibody, horseradish peroxidase (HRP) goat anti-mouse IgG (Zymed), was added at a concentration on 1:2000 in PBS/T/W + 10% FBS. The secondary antibody was detected by addition of a developing solution [0.02% 3,3’,5,5’-tetramethylbenzidine (TMB) and 0.01% hydrogen peroxide in 0.1 M sodium acetate buffer, pH 6.0] and allowing the color to develop for 3 minutes. The reaction was stopped by addition of 1 M phosphoric acid and the absorbance was measured at 450 nm.

### T-cell Proliferation

Spleens were harvested and placed in cold C tubes (MiltenyiBiotec) containing 3 mL of 10 mM phosphate buffered saline (PBS), pH 7.2, 0.5% bovine serum albumin (BSA), 2 mM EDTA. Tubes remained on ice until the spleens were homogenized using the GentleMACSTMDissociator (MiltenyiBiotec). Homogenized spleens were poured through a 40 µm cell strainer (Fisher Biosciences) and the dissociated cells were collected in a 50 ml conical tube. The cell strainer was washed with 5 ml of PBS, pH 7.2, 0.5% BSA, 2 mM EDTA to remove any cells that were attached to the mesh screen. Cells were pelleted by centrifuging the tubes at 500 x g for 7 minutes. Red blood cells (RBC) were lysed by addition of 1 ml of RBC Lysing Buffer (Sigma) and mixing for 2.5 min. The reaction was stopped by addition of 20 mL of RPMI 1640 medium to each tube. Splenocytes were pelleted by centrifugation at 500 x g for 7 minutes. The supernatant was decanted and the cells were resuspended in 5 ml of complete medium (RPMI 1640 with 2 mM L-glutamine, 10% FBS, 100 U/ml penicillin and 100 µg/ml streptomycin). Cell count was determined by adding 5 µL of cells to 95 µL of 0.4% Trypan Blue and using 10 µL of that solution to be counted by the Countess Automated Cell Counter (InvitrogenTM).

For the proliferation assay splenocytes were plated in a Corning Costar 96-well round bottom cell culture plate (Sigma-Aldrich) at a cell density of 2.5 × 10^5^ cells in a final volume of 170 µL. Peptides and positive controls (F1-V recombinant protein) were added in a volume of 30 µL at µg and 2 µg per well, respectively. Plates were incubated at 37 °C in a 5% CO_2_ environment for 72 hours. After 72 hours, 1.0 µCi of ^3^H-thymidine was added to each well. Cultures were incubated for an additional 18 hours before the cells were harvested onto a glass filter mat (Skatron) using a cell harvester.

Filters were placed in a 20-mL scintillation vial and allowed to dry for 24 hours. The next day, 10 mL of Opti-Fluor O (PerkinElmer) scintillation fluid was added to each vial and cell proliferation was determined by measuring the levels of ^3^H-thymidine incorporation using a scintillation counter. The response of a single mouse was considered positive if the stimulation index (SI) was greater than 2, which corresponded to 2 standard deviations above the average proliferation of unstimulated cultures. Peptides were considered immunodominant if six or more mice responded.

### Prediction of MHCII affinity

Peptides having high affinity for the MHCII molecules in each mouse strain were identified using the NetMHCII 2.3 Server (4). This server makes use of a neural network-based approach that is trained on a large data set of more than 14,000 quantitative peptide MHC binding values that cover 14 alleles. Due to the importance of the peptide flanking residues, each peptide encoded into the method includes information about the peptide binding core and the length and composition of the residues flanking the core. The sequences for the V antigen and F1 were input to the web-based server and the IC_50_ values of the 17-mers to I-A^b^, I-A^k^, and I-A^d^ were retrieved.

### Antigen Processing Likelihood

To predict antigen processing likelihood (APL), we use the algorithm described in (36). Our algorithm works by first aggregating input sources of conformational stability data. We typically use four sources of data: sequence entropy, crystallographic b-factors, COREX score, and solvent accessible surface area. The aggregation procedure works by combining z-scores computed from the given sources of data in order to eliminate variations in range and scale. Thus, other sources of data, as well as a different number of sources can be used as input; we can also vary the contribution of each source of data based on its estimated importance. In the resulting stability profile, we consider residues having a positive z-score as “stable” and all other residues as “unstable.” Computing APL proceeds by upweighting all regions in which the stability profile transitions from stable to unstable regions (or vice versa). Upweighting at these regions of transitional stability attempts to capture the increased likelihood of proteolysis, which in turn would capture the corresponding increased likelihood of epitopes adjacent to proteolytic sites.

In this paper we follow the upweighting procedure described in (36), but with generalized constants which are optimized. For clarity we describe the standard approach and then describe the generalization. For a given transitional region, suppose we have *S* residues in the stable component and *U* residues in the unstable component. The upweighting scheme scales up the weights of the *S/3* residues closest to the transition by a magnification factor 2. The weights of *U/2* residues closest to the transition are set in a linearly increasing fashion, starting from the midpoint of the unstable region and ending at the midpoint of the upweighted stable region.

### Parameter Optimization

To improve performance, we generalize the latter algorithm by conducting a search over the optimal values of the constants used in the algorithm. To do this, we maximize positive predictive value (PPV, calculated as the ratio of true positives to identified positives. We optimize parameters for each antigen of interest (the “test antigen”), by utilizing a methodology analogous to the standard cross-fold validation methodology from machine learning. The parameters we consider are the individual weights of the input data sources (i.e., “data parameters”) and the parameters of the upweighting procedure (i.e., “algorithm parameters”) which define the magnification factor, and portions of stable and unstable regions that are upweighted. The range considered for each data parameter was between 0 and 1, in increments of 0.1 with the constraint that they sum to 1. The ranges for algorithm parameters were as follows: loop sizes of 0 to 30 were considered in increments of 5 residues, flank sizes of 9 to 30 were considered in increments of 5 residues. Finally, when applicable the ratio of APL to MHC scores (i.e., for single allele datasets) was searched in the range of 0 to 1 in increments of 0.1.

We perform the optimization of parameters for each test antigen individually, in two stages. We refer to the remaining antigens as the “training set.” First, we perform an optimization for the algorithm parameters on all antigens in the training set (either from C57BL/6 mice or human subjects). We then fix these algorithm parameters and optimize the data parameters in the training set. For the final data parameters we take the average relative weights of the four sources yielding the top 10% PPV values. We then conduct a final round with these data parameters to reoptimize the algorithm parameters to obtain the final choice for the test antigen.

### Statistical tests

The Wilcoxon Signed Rank Test for identification of positive T-cell response, ROC-AUC with significance for prediction accuracy, One-way ANOVA with repeated measures and Tukey multiple comparisons for hit-frequency of prediction methods versus “Random” selection of mouse epitopes (epitope frequency), paired T-test for APL vs. “Random” selection of human epitopes, and paired T-test for number of hits predicted by Combined vs. MHCII were calculated using GraphPad Prism.

## Author Contributions

TC designed and conducted experiments and wrote the manuscript. DM designed and conducted experiments. PB, PM, and NK conducted experiments. AB and RM conducted computational studies and analyzed data. SJL designed experiments, wrote and edited the manuscript.

## Funding

This work was supported by NIH grant R21-AI122199 to R.M. Proteomics work was supported by P20 RR018766, P20 GM103514, P30 GM103514, and a special appropriation from the Louisiana State University School of Medicine Office of the Dean.

### Acknowledgments

We would like to thank N. Kalaya Steede for assistance with animal studies, Jessie Guidry for assistance with proteomics. The following reagents were obtained through BEI Resources, NIAID, NIH: Yersinia pestis F1-V Fusion Protein, Recombinant from Escherichia coli, NR-4524; Yersinia pestis F1-V Fusion Protein, Monomer-Enriched Antigen, Recombinant from Escherichia coli, NR-2561; Yersinia pestis LcrV Protein, Recombinant from Escherichia coli, NR-32875; Peptide Array, Yersinia pestis F1 capsule antigen, NR-2866; Peptide Array, Yersinia pestis V antigen, NR-2867.

